# The Complete Chloroplast Genome of *Tetraselmis desikacharyi* (Chlorodendrophyceae) and Phylogenetic Analysis

**DOI:** 10.1101/591727

**Authors:** Linzhou Li, Haoyuan Li, Sunil Kumar Sahu, Yan Xu, Hongli Wang, Hongping Liang, Sibo Wang

**Affiliations:** School of Biology and Biological Engineering, South China University of Technology, 510006, China; China National GeneBank, BGI-Shenzhen, Jinsha Road, Shenzhen 518120, China; State Key Laboratory of Agricultural Genomics, BGI-Shenzhen, Shenzhen 518083, China

## Abstract

*Tetraselmis desikacharyi* is a marine alga, known as an important plankton for aquaculture as a feed organism. However, the genomic study on this class is rare. Here, we present a complete Chlorodendrophyceae chloroplast genome of *T. desikacharyi*, belonging to Chlorodendrophyceae with a full length of 149,934bp, characterized by a very small single-copy (SSC) region without any genes and a large inverted repeat (IR) region. A maximum-likelihood (ML) phylogenetic analysis was performed using three kinds of data comprising 50 protein-coding genes, which placed the Chlorodendrophyceae as a deep-diverging lineage of the core Chlorophyta.

*T. desikacharyi*, characterized by their intense green colored chloroplast, is important for aquaculture as a feed organism, and for studying plankton growth cycles due to their fast growth rate (Arora et al., 2013; Norris et al., 1980). However, only two chloroplast genomes have been published in the past years for Chlorodendrophyceae (Turmel et al., 2016), which makes the phylogenetic relationship between Chlorodendrophyceae and UTC clade (Ulvophyceae, Trebouxiophyceae, Chlorophyceae) a mystery.

Here, we reported a complete chloroplast genome of *T. desikacharyi*. The samples were collected from marine envionment at Rochigou, Ile de Batz, Bretagne, France (48°44′43″N, 4°00′35″W). The voucher specimen has been deposited in the Culture Collection of Algae at the University of Cologne (Strain number: CCAC 0023) and were grown in Waris-H culture medium (http://www.ccac.uni-koeln.de/) (McFadden and Melkonian, 1986). Total DNA was extracted using a modified CTAB protocol (Sahu et al., 2012). BGISEQ-500 platform was used to generate approximately 50Gb high quality, paired-end (100bp) reads. CLC-Assembly-Cell-5.1.1 (http://www.clcbio.com/) was used to remove the adapter and unpaired reads. The complete chloroplast genome was assembled by the seed-extension-based *de novo* assembler NOVOPlasty2.5.9 (Dierckxsens et al., 2017), based on an rbcL seed gene sequence of *Chlamydomonas reinhardtii*. The assembled chloroplast genome was annotated using a web-based program GeSeq (Tillich et al., 2017), and the protein-coding genes were further annotated by GeneWise v2.4.1 (Birney and Durbin, 2000). The plastid genome has been submitted to the CNGB Nucleotide Sequence Archive (CNSA: https://db.cngb.org/cnsa; accession number: CNP0000390).

The complete plastid genome of *T. desikacharyi* has a total length of 149,934 bp, with a typical quadripartite structure composed of a large single-copy (LSC) region (71,639bp) and an SSC region (1,611bp) interspersed between the IRa/b region (38,343bp). It encodes a total of 81 protein-coding genes, 6 ribosomal RNAs, and 37 transfer RNAs.

To further investigate its phylogenetic position, a maximum likelihood phylogenetic tree was generated using RAxML based on complete chloroplast genome sequences along with 20 other Chlorophyta species. Three different data sets were used for the phylogenetic tree construction: (1) all nucleotide positions; (2) the first and second codon nucleotide positions; (3) the amino acid of 50 protein-coding genes. GTRCAT (PROTGAMMAWAG for AA) model was used with 500 bootstraps. The results revealed that the phylogenetic relationship using the three different data sets were highly consistent at the family level, which places the Chlorodendrophyceae as a deep-diverging lineage of the core Chlorophyta (Figure 1).

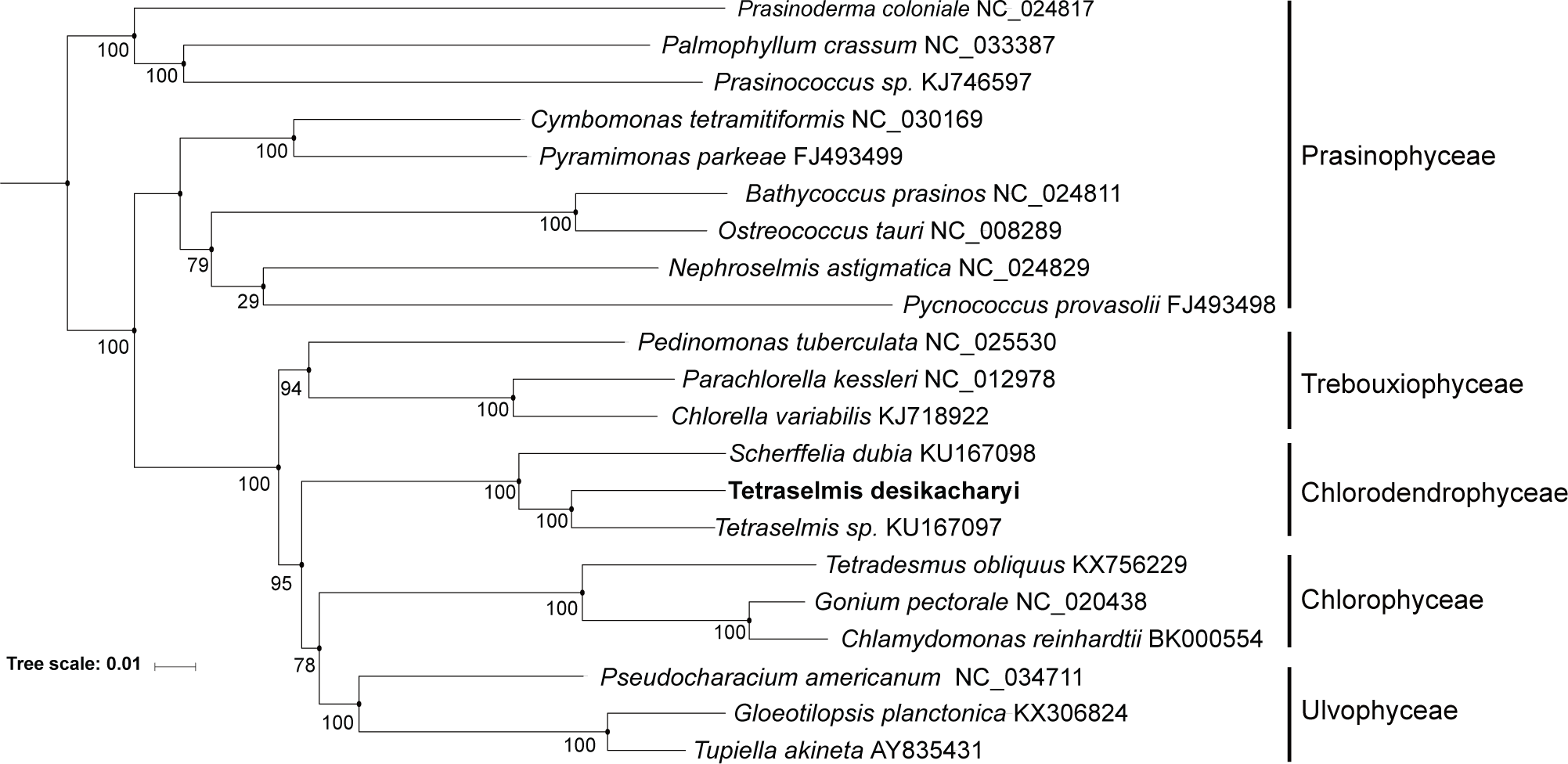

